# Closing coronavirus spike glycoproteins by structure-guided design

**DOI:** 10.1101/2020.06.03.129817

**Authors:** Matthew McCallum, Alexandra C. Walls, Davide Corti, David Veesler

**Affiliations:** Department of Biochemistry, University of Washington, Seattle, Washington 98195, USA; Humabs Biomed SA, a subsidiary of Vir Biotechnology, 6500 Bellinzona, Switzerland

## Abstract

The recent spillover of SARS-CoV-2 in the human population resulted in the ongoing COVID-19 pandemic which has already caused 4.9 million infections and more than 326,000 fatalities. To initiate infection the SARS-CoV-2 spike (S) glycoprotein promotes attachment to the host cell surface, determining host and tissue tropism, and fusion of the viral and host membranes. Although SARS-CoV- 2 S is the main target of neutralizing antibodies and the focus of vaccine design, its stability and conformational dynamics are limiting factors for developing countermeasures against this virus. We report here the design of a prefusion SARS-CoV-2 S ectodomain trimer construct covalently stabilized in the closed conformation. Structural and antigenicity analysis showed we successfully shut S in the closed state without otherwise altering its architecture. Finally, we show that this engineering strategy is applicable to other β-coronavirus S glycoproteins and might become an important tool for vaccine design, structural biology, serology and immunology studies.

---

In the past two decades, three zoonotic coronaviruses crossed the species barrier to cause severe pneumonia in humans: (i) severe acute respiratory syndrome coronavirus (SARS-CoV), that was associated with an epidemic in 2002-2003 and a few additional cases in 2004^1,2^, (ii) Middle-East respiratory syndrome coronavirus (MERS-CoV), which is currently circulating in the Arabian peninsula^3^, and (iii) SARS-CoV-2, the etiological agent of the ongoing COVID-19 pandemic^4,5^. SARS-CoV-2, was discovered in December 2019 in Wuhan, Hubei Province of China, was sequenced and isolated by January 2020^4,6^ and has infected over 4.9 million people with more than 326,000 fatalities as of May 20^th^ 2020. No vaccines or specific therapeutics are licensed to treat or prevent infections from any of the seven human-infecting coronaviruses with the exception of Remdesivir^7,8^ which was recently approved by the Food and Drug Administration for emergency use for COVID-19 treatment.

Coronaviruses gain access to host cells using the homotrimeric transmembrane spike (S) glycoprotein protruding from the viral surface^9^. S comprises two functional subunits: S_1_ (encompassing the A, B, C and D domains) and S_2_. These subunits are responsible for binding to the host cell receptor and fusion of the viral and cellular membranes, respectively^10^. For many coronaviruses, including the newly emerged SARS-CoV-2, S is cleaved at the boundary between the S_1_ and S_2_ subunits which remain non-covalently bound in the prefusion conformation^10-18^. The distal S_1_ subunit comprises the receptor-binding domain(s), and contributes to stabilization of the prefusion state of the membrane-anchored S_2_ subunit which contains the fusion machinery^10,17,19-25^. For all coronaviruses, upon receptor binding S is further cleaved by host proteases at the S_2_’ site located immediately upstream of the fusion peptide^14,16,26^. This cleavage has been proposed to activate the protein for membrane fusion *via* extensive irreversible conformational changes^13-16,19,27,28^. As a result, coronavirus entry into susceptible cells is a complex process that requires the concerted action of receptor-binding and proteolytic processing of the S protein to promote virus-cell fusion.

Viral fusion proteins, including coronavirus S glycoproteins, fold in a high-energy, kinetically-trapped prefusion conformation found at the viral surface before host cell invasion^29^. This metastable state is activated with exquisite spatial and temporal precision upon encounter of a target host cell by one or multiple stimuli such as pH change^30,31^, proteolytic activation^13,15^ or protein-protein interactions^32^. The ensuing irreversible and large-scale structural changes of viral fusion proteins are coupled to fusion of the viral and host membrane to initiate infection. As a result, the postfusion state of a viral fusion protein is the lowest energy conformation (i.e. ground state) observed throughout the reaction coordinates^29^. A notable exception to this general pathway is the vesicular stomatitis virus fusion glycoprotein G that can reversibly fold from the postfusion to the prefusion conformation^31,33,34^.

The intrinsic metastability of viral fusion proteins – which is oftentimes magnified by working with ectodomain constructs lacking the transmembrane and cytoplasmic segments – has posed challenges for studying the structure and function of these glycoproteins and for vaccine design. As a result, a variety of approaches have been implemented to stabilize these fragile glycoproteins. Proline substitutions preventing refolding to an elongated *α*-helical structure observed in postfusion influenza virus hemagglutinin were reported as a promising strategy to stabilize the prefusion state of this widely studied viral glycoprotein^35^. Engineering approaches based on this concept along with introduction of designed disulfide bonds and other mutations have subsequently been utilized for stabilizing the prefusion conformation of other class I fusion proteins, such as the SOSIP mutations in the HIV-1 envelope glycoprotein^36-39^. Structure-guided prefusion stabilization via introduction of disulfide bonds and cavity-filling mutations was successfully implemented for the respiratory syncytial virus fusion glycoprotein^40^ (DS-Cav1) and parainfluenza virus 1-4 fusion glycoproteins^41^. Designed disulfide bonds have also proven useful to enhance the prefusion stability of the Hendra virus fusion glycoprotein^42^, mutations which were later applied to the Nipah virus fusion protein^43^. Finally, the introduction of double proline substitutions, herein 2P, to prevent fusogenic conformational changes of MERS-CoV S^20^ and SARS-CoV S^44^ was shown to stabilize the prefusion states of these glycoproteins. These results provided proof-of-concept of the broad applicability of this approach to coronavirus S glycoproteins, which was subsequently confirmed by its successful use for SARS-CoV-2 S structural studies^18,45,46^. In spite of these advances, the conformational dynamics and limited stability of the SARS-CoV-2, SARS-CoV and MERS-CoV S glycoproteins remain a challenge that needs to be overcome to accelerate structural studies of the immune response elicited by coronavirus infections and vaccine design. Recent reports of the observation of postfusion trimers at the surface of purified authentic SARS-CoV-2^47^ and of spontaneous refolding of a fraction of S trimers upon detergent-solubilization^48^ showcase these limitations.

We report here the design of a prefusion-stabilized SARS-CoV-2 S ectodomain trimer construct engineered to remain in the closed conformation through introduction of an intermolecular disulfide bond. Single-particle cryo-electron microscopy analysis of this glycoprotein coupled with ELISA assays unambiguously demonstrated that our strategy successfully shut S in the closed state without otherwise altering its architecture, as evaluated by binding to a panel of human monoclonal neutralizing antibodies and a COVID-19 convalescent plasma. We show that this covalent stabilization strategy is applicable to other *β*-coronavirus S glycoproteins and envision it might become an important tool for vaccine design, structural biology, serology and immunology studies.

## Structure-based design of a SARS-CoV-2 S trimer stabilized in the closed conformation

Structural fluctuations of the receptor-binding S^B^ domain (also known as RBD), from a closed to an open conformation, enables exposure of the receptor-binding motif which mediates interaction with angiotensin-converting enzyme 2 (ACE2) for SARS-CoV- 2^6,18,49-54^ and SARS-CoV^55,56^, or dipeptidyl-peptidase 4 for MERS-CoV^57,58^ **(Fig 1 a-b)**. Receptor-engagement or interaction with the Fab fragment of the S230 neutralizing monoclonal antibody were previously shown to induce the SARS-CoV S cascade of conformational changes leading to membrane fusion, which we proposed to proceed through a molecular ratcheting mechanism^22,59^. Although S^B^ opening and/or displacement/shedding of the S_1_ subunit is expected to occur for all coronavirus S glycoproteins, to free the S_2_ subunit during membrane fusion^60^, such changes have only been observed for SARS-CoV-2, SARS-CoV and MERS-CoV S glycoproteins^20-23,44,59^. Since these three S trimers suffer from limited stability compared to S trimers that have not been observed to open, such as HCoV-OC43 S^24^, we reasoned that arresting the first step of SARS-CoV-2 S refolding toward the postfusion state might enhance the stability of the prefusion state.

**Figure 1.**
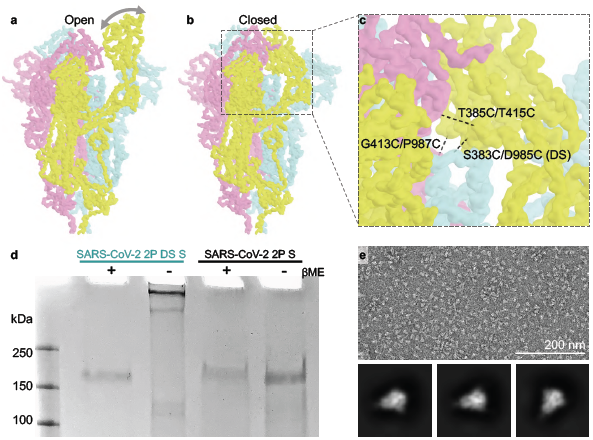
Structure-based engineering of a SARS-CoV-2 S trimer in the closed conformation. **a-b**, CryoEM structures of SARS-CoV-2 S with one S^B^ receptor-binding domain open (a, PDB 6VYB) and in the closed state (b, PDB 6VXX) used as a basis for the design of intermolecular disulfide bonds^18^. **c**, Pairs of residues mutated to create potential disulfide bonds are shown with dashed black lines between the alpha-carbons. In panels a-c, each S protomer is colored distinctly. **d**, SDS-PAGE analysis in reducing and non-reducing conditions showing formation of an intermolecular disulfide bond. *β*ME: *β*-mercapto-ethanol. **e**, Electron micrograph of negatively stained SARS-CoV-2 2P DS S confirming proper folding of the designed protein construct (top) and representative 2D class averages (bottom).

Based on these considerations, we set out to engineer SARS-CoV-2 S stalled in the closed conformation of the three S^B^ receptor-binding domains through introduction of disulfide bonds within the ectodomain trimer construct we previously used to determine structures of the closed and open conformations^18,46^. Specifically, our construct harbored an abrogated furin S_1_/S_2_ cleavage site (R682S, R683G and R685G)^10,24,59,61^, two consecutive proline stabilizing mutations (K986P and V987P)^20,44^ and a C-terminal foldon trimerization domain^62^. We designed the following pairs of cysteine substitutions aimed at introducing three inter-molecular disulfide bonds per trimer: S383C/D985C, G413C/P987C, and T385C/T415C **(Fig 1c)**. Out of the three pairs of substitutions tested, only the S383C/D985C (termed SARS-CoV-2 2P DS S) could be recombinantly expressed using HEK293 Freestyle cells and purified. SDS-PAGE analysis of SARS-CoV-2 2P DS S in reducing and non-reducing conditions demonstrated that the engineered disulfide bond was indeed correctly introduced **(Fig 1d)**. Further characterization of purified SARS-CoV-2 2P DS S using negative staining electron microscopy indicated proper homotrimer folding and assembly **(Fig 1e)**.

## CryoEM structure of a prefusion SARS-CoV-2 S trimer stabilized in the closed conformation

Since the DS substitutions connect regions of the S glycoprotein that are far apart upon S^B^ receptor-binding domain opening and transition to the postfusion S state^60,63^, we expected this protein construct to be trapped in the closed S state via molecular stapling. To validate our design strategy, we used single particle cryo-electron microscopy to analyze the conformational landscape of SARS-CoV-2 2P DS S **(Table 1 and Fig S1)**. 3D classification of the cryoEM dataset demonstrated that all particle images clustered in 3D reconstructions of the closed S trimer **(Fig S2)**. In contrast, about half of the particle images selected from our previous SARS-CoV-2 2P S apo dataset corresponded to the closed S trimer whereas the other half was accounted for by a partially open S trimer^18^. These results therefore indicate we successfully engineered a shut closed S trimer.

**Table 1.**
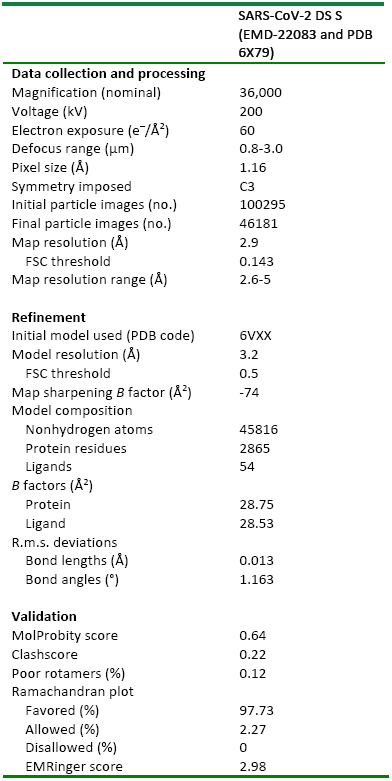
Cryo-EM data collection, refinement and validation statistics.

|  |
| --- |
| SARS-CoV-2 DS S<br>(EMD-22083 and PDB<br>6X79) |

**Data collection and processing**
|  |  |
| --- | --- |
| Magnification (nominal) | 36,000 |
| Voltage (kV) | 200 |
| Electron exposure (e<sup>-</sup>/Å<sup>2</sup>) | 60 |
| Defocus range (μm) | 0.8-3.0 |
| Pixel size (Å) | 1.16 |
| Symmetry imposed | C3 |
| Initial particle images (no.) | 100295 |
| Final particle images (no.) | 46181 |
| Map resolution (Å) | 2.9 |
| FSC threshold | 0.143 |
| Map resolution range (Å) | 2.6-5 |

**Refinement**
|  |  |
| --- | --- |
| Initial model used (PDB code) | 6VXX |
| Model resolution (Å) | 3.2 |
| FSC threshold | 0.5 |
| Map sharpening *B* factor (Å<sup>2</sup>) | -74 |
| Model composition |  |
| Nonhydrogen atoms | 45816 |
| Protein residues | 2865 |
| Ligands | 54 |
| *B* factors (Å<sup>2</sup>) |  |
| Protein | 28.75 |
| Ligand | 28.53 |
| R.m.s. deviations |  |
| Bond lengths (Å) | 0.013 |
| Bond angles (°) | 1.163 |

**Validation**
|  |  |
| --- | --- |
| MolProbity score | 0.64 |
| Clashscore | 0.22 |
| Poor rotamers (%) | 0.12 |
| Ramachandran plot |  |
| Favored (%) | 97.73 |
| Allowed (%) | 2.27 |
| Disallowed (%) | 0 |
| EMRinger score | 2.98 |

We subsequently determined a 3D reconstruction of SARS-CoV-2 2P DS S at 2.9 Å resolution (applying 3-fold symmetry) **(Fig 2 a-b)**. As the SARS-CoV-2 2P DS S structure was obtained with much fewer particles (46,181 particles) than all other studies to date^18,45,46^, we speculate that this engineered construct has enhanced stability relative to the construct lacking the DS mutations. The cryoEM map shows a good agreement with our previously determined structure of SARS-CoV-2 2P S in the closed conformation^18^ with which it could be superimposed with a C*α* root mean square deviation of 1.36 Å over 959 aligned alpha-carbons **(Fig 2 c)**. The cryoEM density also resolves the disulfide bond between a S^B^ receptor-binding domain residue facing towards the fusion machinery (S383C) and the hairpin preceding the S_2_ subunit central helix (D985C) from a neighboring protomer, the latter residue being located directly upstream the K986P and V987P prefusion-stabilizing mutations **(Fig 2 d)**. These findings not only validate the structure-based design strategy but also show that it did not induce distortions of the S trimer. We note that density at the C-terminal stem helix was not resolved in SARS-CoV- 2 2P DS S – accounting for 6 amino acid residues – whereas this region was visible in previously reported apo S maps^18,45^ but not in the SARS-CoV-2 S/S309 neutralizing antibody complex map^46^.

**Figure 2.**
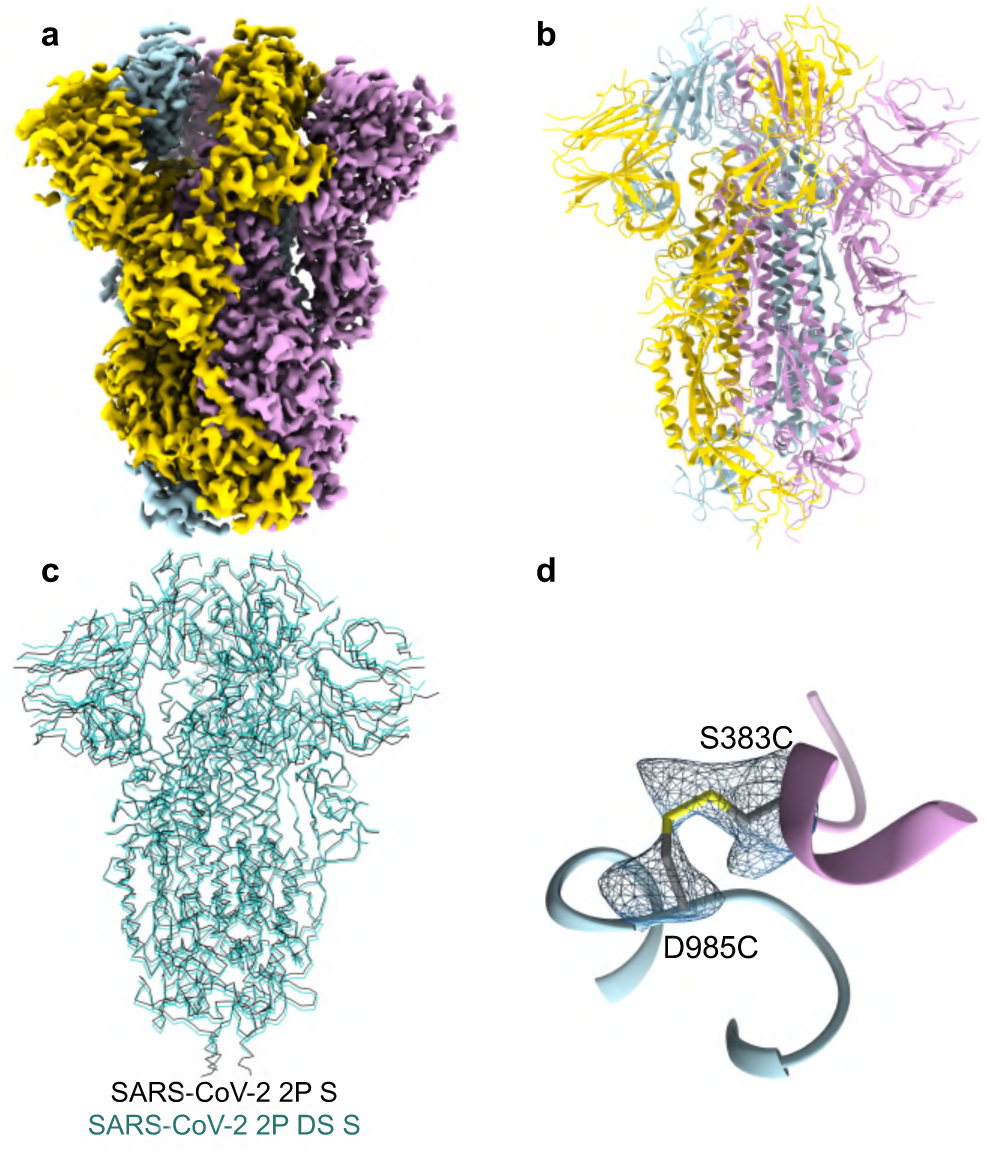
CryoEM structure of the closed SARS-CoV-2 DS S glycoprotein. **a**, CryoEM map of the SARS-CoV-2 DS S trimer in the closed conformation at 2.9Å resolution. **b**, Ribbon diagram of the SARS-CoV-2 DS S trimer atomic model in the same orientation as in panel a. In panels a-b, each S protomer is colored distinctly. **c**, Superimposition of the SARS-CoV-2 DS S trimer (green) to the coordinates from the 2.8 Å SARS-CoV-2 S structure in the closed conformation, PDB 6VXX^18^ (black). **e**, Zoomed-in view of the designed disulfide bond with the corresponding region of cryoEM density shown as a blue mesh.

## Evaluation of SARS-CoV-2 2P DS antigenicity

The high structural similarity between the SARS-CoV-2 2P DS S structure presented here and our previously reported SARS-CoV-2 2P S structure (in the closed conformation)^18^, led us to hypothesize that they would have similar antigenicity profiles – besides the lost ability to interact with binders requiring opening of the S^B^ receptor-binding domains (e.g. the receptor-binding motif) for the former construct. To probe the influence of the introduced disulfide bond on antigenicity, we evaluated binding of SARS-CoV-2 2P DS S and SARS-CoV-2 2P S, side-by-side, to a panel of human neutralizing antibodies by ELISA. Binding to S309 was indistinguishable between the two constructs **(Fig. 3 a)**, in agreement with the fact this antibody recognizes an epitope within the S^B^ receptor-binding domain that remains accessible in both the open and closed states^46^. Furthermore, we observed a dose-dependent response for binding of the receptor-binding motif-targeted S2H14 antibody to SARS-CoV-2 2P S which was dampened by several orders of magnitude with SARS-CoV-2 2P DS S, as expected due to conformational masking of the receptor-binding motif in the closed conformation^18^ **(Fig. 3 b)**. Although the S304 antibody interacted with SARS-CoV-2 2P S in a concentration-dependent manner, it did not bind to SARS-CoV-2 2P DS S **(Fig. 3 c)**. Since we previously demonstrated that S304 recognizes an epitope distinct from both the receptor-binding motif and the S309 epitope, it is expected that this antibody binds to a cryptic epitope that is only accessible upon S^B^ opening, as is the case for the CR3022 antibody^64-67^. Collectively, these findings validate that SARS-CoV-2 2P DS S is in a native, closed conformation and illustrate the usefulness of this protein construct to investigate epitopes recognized by neutralizing antibodies.

**Figure 3.**
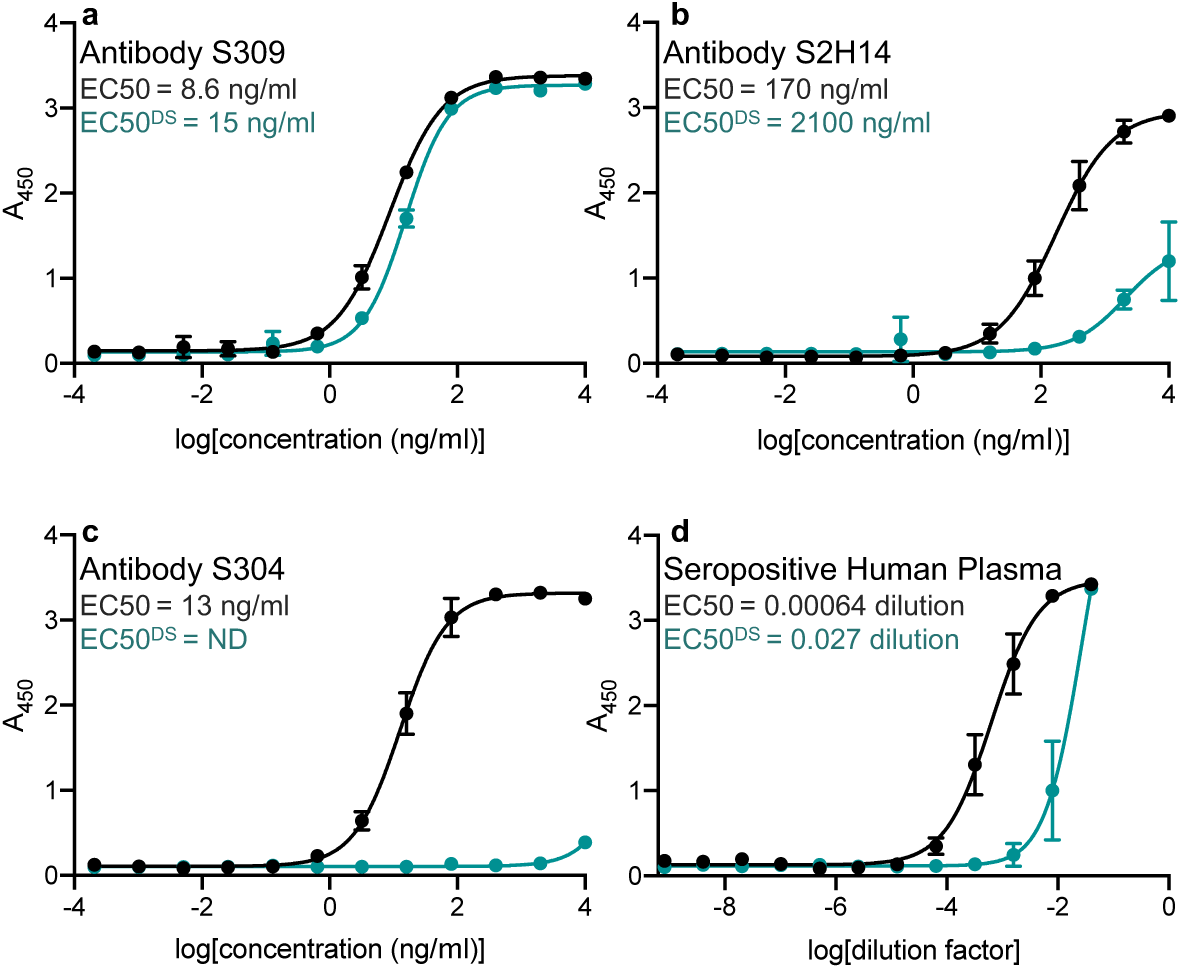
Evaluation of SARS-CoV-2 2P DS S antigenicity. **a-c**, Binding of serially diluted concentrations of the human neutralizing antibodies S309 (a), S2H14 (b) and S304 to immobilized SARS-CoV-2 2P DS S (green) or SARS-CoV-2 2P S (black). **d**, Binding of a serial dilution of a potent neutralizing convalescent plasma to immobilized SARS-CoV-2 2P DS S (green) or SARS-CoV-2 2P S (black). n=2 experiments (technical replicates).

We subsequently assessed binding of a COVID-19 convalescent patient plasma sample to SARS-CoV-2 2P DS S and SARS-CoV-2 2P S by ELISA. The sample was obtained from a Washington State donor who was selected based on its unusually high plasma antibody neutralization titer (mean half-maximal inhibition reciprocal dilution of ∼1/800, **Fig S3**). Comparison of half-maximal binding titers showed that recognition of SARS-CoV-2 2P DS S was ∼40-fold weaker than that of SARS-CoV-2 2P S **(Fig. 3 d)**. This difference likely reflect the proportion of antibodies directed to the receptor-binding motif or cryptic epitopes similar to the one recognized by S304 in this plasma sample. As SARS-CoV-2 2P DS S only displays closed S^B^ receptor-binding domains within the context of a folded trimer, we suggest it will be a useful tool for serology studies aiming at evaluating antibody responses in COVID-19 patients, and could complement tests using ACE2 inhibition as a proxy for evaluating the presence of neutralizing antibody titers in the human population.

## Disulfide stapling is a broadly applicable design strategy for coronavirus S glycoproteins

Next, we set out to test the general applicability of the DS stabilizing strategy identified for SARS-CoV-2 S to other coronavirus S glycoproteins. Based on the high sequence and structural conservation of the residues involved in – and adjacent to – the engineered disulfide bond **(Fig. 4 a)**, we hypothesized that it might be transferable to SARS-CoV S, which shares 80% sequence identity with SARS-CoV-2 S (lineage B).

**Figure 4.**
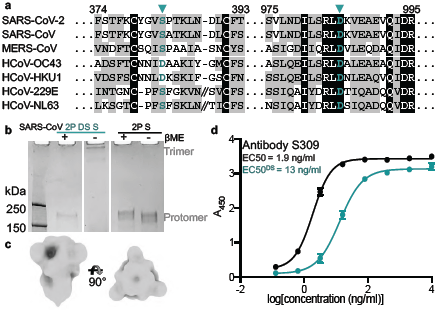
Design of SARS-CoV 2P DS S. **a**, Sequence alignment showing the conservation of the residues involved in and adjacent to the designed disulfide bond across SARS-CoV and SARS-CoV-2 S glycoproteins. Residues are highlighted if they are identical in the alignment (black) or conservatively substituted (grey). Residues are numbered according to the SARS-CoV-2 S sequence. Green triangles highlight substituted residues. **b**, SDS-PAGE analysis in reducing and non-reducing conditions showing formation of an intermolecular disulfide bond. *β*ME: *β*-mercapto-ethanol. **c**, 3D reconstruction in two orthogonal orientations of negatively stained SARS-CoV 2P DS S confirming proper folding of the designed protein construct. **d**, Binding of various concentrations of the human neutralizing antibody S309 to immobilized SARS-CoV 2P DS S (green) or SARS-CoV 2P S (black). n=2 experiments (technical replicates).

SDS-PAGE analysis of SARS-CoV 2P DS S (S370C/D969C mutants) glycoprotein recombinantly expressed using HEK293F cells confirmed that it formed a high-molecular weight species in non-reducing conditions, consistent with covalent formation of S homotrimers **(Fig. 4 b)**. Likewise, the electrophoretic mobility corresponds to individual S protomers in presence of *β*-mercaptoethanol. Electron microscopy analysis of negatively stained SARS-CoV 2P DS S demonstrated the engineered mutant folded as a globular, closed homotrimer **(Fig. 4 c)**. Finally, we observed dose-dependent binding of the S309 neutralizing antibody to SARS-CoV 2P DS S and SARS-CoV 2P S (albeit stronger to the later protein) validating retention of antigenicity of the designed construct **(Fig. 4 d)**.

## Discussion

Viral glycoprotein engineering is an active field of research fueling vaccine design strategies to elicit potent and/or broad protection against a range of emerging or endemic pathogens. Stabilization of the respiratory syncytial virus fusion glycoprotein in its prefusion conformation (DS-Cav1)^40^ and subsequent fusion to a computationally designed trimeric protein (I53-50A)^68^ are recent breakthroughs illustrating the power of structure-based vaccine design. Ds-Cav1 is currently evaluated in a phase I randomized, open-label clinical trial to assess its safety, tolerability and immunogenicity in healthy adults (NCT03049488). Furthermore, multivalently-displayed Ds-Cav1 genetically fused to the I53-50 nanoparticle has been shown to further improve elicitation of high titers of neutralizing antibodies^68^. Similarly, the development of HIV-1 SOSIP constructs have revolutionized the field of HIV-1 structural vaccinology and immunology^36^.

The recent emergence of SARS-CoV-2, the virus responsible for the ongoing COVID-19 pandemic, showcases the urgent need to explore strategies to expedite coronavirus vaccines and therapeutics design initiatives as well as structural and serology studies. Prefusion-stabilization of MERS-CoV S through the aforementioned introduction of two proline substitutions was previously shown to elicit enhanced neutralizing antibody titers against multiple MERS-CoV isolates in mice^20^. However, the limited stability and conformational dynamics of the SARS-CoV-2 2P S, SARS-CoV 2P S and MERS-CoV 2P S ectodomain trimers indicate that further improvements are needed to increase their shelf life and/or to manipulate their conformational states.

We report here a strategy to produce prefusion-stabilized, closed coronavirus ectodomain trimers and show it is broadly applicable to at least the *β*-genus which accounts for 5 out of 7 human-infecting coronaviruses, including the most pathogenic members (i.e. SARS-CoV-2, SARS-CoV and MERS-CoV). By symmetrizing and stabilizing S proteins, we expect the DS mutation be a useful tool for the research community, enabling high-resolution structural studies of antibody complexes, and for characterizing the humoral immune response in infected or vaccinated individuals and animals.

We hypothesize that the design strategy described here might improve the breadth of neutralizing antibodies elicited via masking of the highly immunogenic receptor-binding motif. The tradeoff, however, will be dampening of induction of antibodies targeting the receptor-binding motif which is typically recognized by potent neutralizing antibodies but with narrow breadth between coronaviruses due to poor conservation. For the same reasons, we also envision that S glycoprotein constructs shut in the closed conformation could assist in isolating broadly neutralizing antibodies effective against multiple viruses belonging to distinct (sub)genera.

Finally, as demonstrated here, comparing the reactivity of DS constructs with protein constructs exhibiting the full range of S^B^ receptor-binding domain conformations will allow to evaluate the fraction of antibodies recognizing the receptor-binding motif and/or cryptic epitopes in serology studies to provide a detailed understanding of the humoral immune response elicited upon infection or vaccination. Given that receptor-binding motif targeting antibodies are typically neutralizing, this comparison may serve as a proxy for estimating whether a patient has neutralizing antibodies or not.

## Acknowledgements

This study was supported by the National Institute of General Medical Sciences (R01GM120553 to D.V.), the National Institute of Allergy and Infectious Diseases (HHSN272201700059C to DV, R01AI141707 and R01AI140891 to J.D.B), a Pew Biomedical Scholars Award (D.V.), an Investigators in the Pathogenesis of Infectious Disease Award from the Burroughs Wellcome Fund (D.V. and J.D.B.), the Bill & Melinda Gates Foundation (OPP1156262 to D.V.), Fast Grants (to D.V.) and the University of Washington Arnold and Mabel Beckman cryoEM center. We are grateful to Jesse Bloom, Adam Dingens and Helen Chu for providing the COVID-19 convalescent plasma sample and to the study participant who contributed this plasma.

## Author contributions

M.M., A.C.W. and D.V. designed the experiments. A.C.W. and M.A.T. expressed and purified the proteins. M.M. carried out ELISAs. M.M prepared samples for cryoEM, collected and processed the data. M.M and D.V built and refined the atomic model. D.C. contributed key reagents. M.M., A.C.W. and D.V. analyzed the data. M.M. and D.V. and prepared the manuscript with input from all authors

## Data availability

The cryoEM map and atomic model have been deposited to the EMDB and PDB with accession numbers EMD-22083 and PDB 6×79.

## MATERIALS AND METHODS

### Design of disulfide mutants

Disulfide mutants were designed using the Disulfide by Design 2 software^69^ and synthetic genes ordered from Genscript.

### Recombinant S ectodomains production

All ectodomains were produced in 500mL cultures of HEK293F cells grown in suspension using FreeStyle 293 expression medium (Life technologies) at 37°C in a humidified 8% CO2 incubator rotating at 130 r.p.m., as previously reported^18^. The culture was transfected using 293fectin (ThermoFisher Scientific) with cells grown to a density of 10^6^ cells per mL and cultivated for three days. The supernatant was harvested and cells were resuspended for another three days, yielding two harvests. Clarified supernatants were purified using a 5 mL Cobalt or Nickel affinity column (Takara). Purified protein was concentrated, and flash frozen in a buffer containing 50 mM Tris pH 8.0 and 150 mM NaCl prior to cryoEM analysis.

### Antibody expression

Recombinant antibodies were expressed in ExpiCHO cells transiently co-transfected with plasmids expressing the heavy and light chain, as previously described^46^.

### Serum preparation

A de-identified COVID-19 patient plasma sample was collected and heat-inactivated at 56 °C for one hour.

### ELISA

20µl of ectodomains (stabilized prefusion trimer) of SARS-CoV-2 or SARS-CoV, or the disulphide stabilized SARS-CoV-2 or SARS-CoV, were coated on 384 well ELISA plates at 1 ng/µl for 16 hours at 4°C. Plates were washed with a 405 TS Microplate Washer (BioTek Instruments) then blocked with 80 µl SuperBlock (PBS) Blocking Buffer (Thermo Scientific) for 1 hour at 37°C. Plates were then washed and 30 µl antibodies were added to the plates at concentrations between 4 × 10^−8^ and 10 ng/µl and incubated for 1 h at 37°C. Plates were washed and then incubated with 30 µl of 1/5000 diluted goat anti-human Fc IgG-HRP (invitrogen A18817). Plates were washed and then 30 µl Substrate TMB microwell peroxidase (Seracare 5120-0083) was added for 5 min at room temperature. The colorimetric reaction was stopped by addition of 30 µl of 1 N HCl. A_450_ was read on a Varioskan Lux plate reader (Thermo Scientific).

### Pseudovirus neutralization assays

Murine leukemia virus (MLV)-based SARS-CoV-2 S-pseudotyped viruses were prepared as previously described^18^. HEK293T cells were co-transfected with a SARS-CoV-2 S encoding-plasmid, an MLV Gag-Pol packaging construct and the MLV transfer vector encoding a luciferase reporter using the Lipofectamine 2000 transfection reagent (Life Technologies) according to the manufacturer’s instructions. Cells were incubated for 5 hours at 37°C with 8% CO_2_ with OPTIMEM transfection medium. DMEM containing 10% FBS was added for 72 hours.

HEK293T cells stably expressing human ACE2^70^ were cultured in DMEM containing 10% FBS, 1% PenStrep and plated into 96 well plates for 16-24 hours. Concentrated pseudovirus with or without serial dilution of COVID-19 convalescent plasma was incubated for 1 hour and then added to the wells after washing 3X with DMEM. After 2-3 hours DMEM containing 20% FBS and 2% PenStrep was added to the cells for 48 hours. Following 48 hours of infection, One-Glo-EX (Promega) was added to the cells and incubated in the dark for 5-10 minutes prior to reading on a Varioskan LUX plate reader (ThermoFisher). Measurements were done in duplicate and relative luciferase units (RLU) were converted to percent neutralization and plotted with a non-linear regression curve fit in PRISM.

### Negative-stain EM sample preparation

All constructs in this study were negatively stained at a final concentration of 0.06 mg/mL using Gilder Grids overlaid with a thin layer of carbon and 2% uranyl formate. Data were acquired using the Leginon software^71^ to control a Tecnai T12 transmission electron microscope operated at 120 kV and equipped with a Gatan 4K Ultrascan CCD detector. The dose rate was adjusted to 50 electrons / Å^2^ and each micrograph was acquired in 1 second. ∼100 micrographs were collected in a single session with a defocus range comprised between −1.0 and −2.5 μm. Data were subsequently processed using cryoSPARC^72^.

### CryoEM sample preparation and data collection

3 µL of SARS-CoV-2 2P DS S at 0.5 mg/mL was applied onto a freshly glow discharged 2.0/2.0 UltraFoil grid (200 mesh). Plunge freezing used a vitrobot MarkIV (ThermoFisher Scientific) using a blot force of 0 and 6.5 second blot time at 100% humidity and 23°C. Data were acquired using the Leginon software^71^ to control a Glacios transmission electron microscope operated at 200 kV and equipped with a Gatan K2 Summit direct detector. The dose rate was adjusted to 8 counts/pixel/s, and each movie was acquired in 50 frames of 200 ms with a pixel size of 1.16 Å at the specimen level. ∼600 micrographs were collected in a single session with a defocus range comprised between −0.8 and −3.0 μm.

### CryoEM data processing

Movie frame alignment, estimation of the microscope contrast-transfer function parameters, particle picking and extraction (with a box size of 352 pixels^2^) were carried out using Warp^73^. Reference-free 2D classification was performed using cryoSPARC^72^ to select well-defined particle images. 3D classification with 50 iterations each (angular sampling 7.5° for 25 iterations and 1.8° with local search for 25 iterations) were carried out using Relion^74^ without imposing symmetry to separate distinct SARS-CoV-2 S conformations. 3D refinements were carried out using non-uniform refinement along with per-particle defocus refinement in cryoSPARC^72^. Particle images were subjected to Bayesian polishing^75^ before performing another round of non-uniform refinement in cryoSPARC^72^ followed by per-particle defocus refinement and again non-uniform refinement. Reported resolutions are based on the gold-standard Fourier shell correlation (FSC) of 0.143 criterion and Fourier shell correlation curves were corrected for the effects of soft masking by high-resolution noise substitution^76^.

### CryoEM model building and analysis

UCSF Chimera^77^ and Coot were used to fit an atomic model (PDB 6VXX) into the cryoEM map. The model was then refined into the map using Rosetta^78-80^ and analyzed using MolProbity^81^, EMringer^82^, and Phenix^83^. Figures were generated using UCSF ChimeraX^84^ and UCSF Chimera^77^.

---

**Supplemental Fig. 1:**
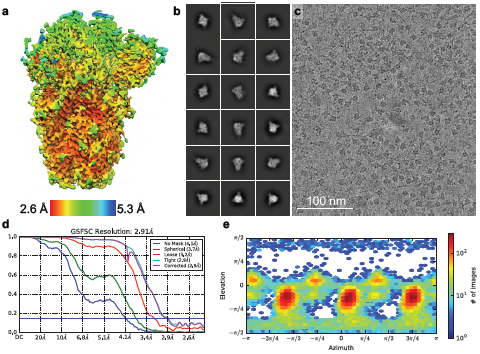
CryoEM data processing and validation. **a**. Local resolution map calculated using cryoSPARC. **b-c**. Representative electron micrograph (c) and class averages (b) of SARS-CoV-2 2P DS S embedded in vitreous ice. Scale bar: 100nm. **d**. Gold-standard Fourier shell correlation curves. The 0.143 cutoff is indicated by horizontal blue line. **e**. Particle orientation distribution plot.

**Supplemental Fig. 2:**
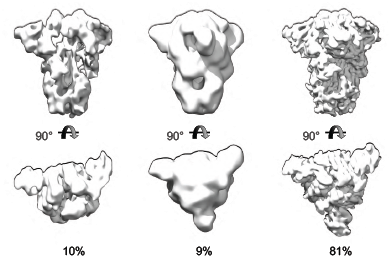
Orthogonal views of the classes obtained by 3D classification. Percentages reflect the proportion of particles assigned to each map.

**Supplemental Fig. 3:**
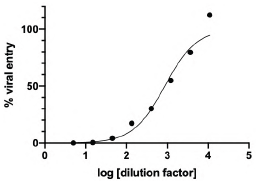
Neutralization of SARS-CoV-2 pseudovirus with human sera from a COVID-19 positive patient.

